# Stat3-mediated Atg7 expression enhances anti-tumor immunity in melanoma

**DOI:** 10.1101/2024.06.10.598284

**Authors:** Sarah M. Zimmerman, Erin Suh, Sofia R. Smith, George P. Souroullas

## Abstract

Epigenetic modifications to DNA and chromatin control oncogenic and tumor suppressive mechanisms in melanoma. EZH2, the catalytic component of the Polycomb repressive complex 2 (PRC2), which mediates methylation of lysine 27 on histone 3 (H3K27me3), can regulate both melanoma initiation and progression. We previously found that mutant *Ezh2*^Y641F^ interacts with the immune regulator Stat3 and together they affect anti-tumor immunity. However, given the numerous downstream targets and pathways affected by EZH2, many mechanisms that determine its oncogenic activity remain largely unexplored. Using genetically engineered mouse models we further investigated the role of pathways downstream of EZH2 in melanoma carcinogenesis and identified significant enrichment in several autophagy signatures, along with increased expression of autophagy regulators, such as Atg7. In this study, we investigated the effect of Atg7 on melanoma growth and tumor immunity within the context of an *Ezh2*^Y641F^ epigenetic state. We found that expression of Atg7 is largely dependent on Stat3 expression and that deletion of Atg7 slows down melanoma cell growth *in vivo*, but not *in vitro*. Atg7 deletion also results in increased CD8+ T cells and reduced myelosuppressive cell infiltration in the tumor microenvironment, suggesting a strong immune system contribution in the role of Atg7 in melanoma progression. These findings highlight the complex interplay between genetic mutations, epigenetic regulators, and autophagy in shaping tumor immunity in melanoma.

## INTRODUCTION

Epigenetic alterations contribute to oncogenesis through multiple mechanisms, from repression of tumor suppressor genes or activation of oncogenes to tumor cell extrinsic mechanisms such as angiogenesis, invasion and anti-tumor immunity ^1–4^. Epigenetic regulators have thus become effective therapeutic targets in multiple solid tumors. One epigenetic complex that is frequently mutated in many solid tumors and directly implicated in antitumor immunity is the Polycomb Repressive Complex 2 and particularly its enzymatic domain, EZH2 ^5,6^. EZH2 possesses histone methyltransferase activity and mediates methylation of histone 3 on lysine 27 (H3K27me). Genetic alterations in EZH2 include both loss- and gain-of-function events, and it can function both as a tumor suppressor ^7–11^ and as an oncogene ^12–16^. A unique point mutation in the methyltransferase domain of EZH2 (SET domain) at tyrosine 641 (Y641), alters its methyltransferase activity and may confer neomorphic functions by promoting unconventional changes to the distribution of H3K27me3 across the genome ^12,17^, with complicated effects on gene expression.

In previous studies, using a genetically engineered mouse model, we found that expression of mutant *Ezh2*^Y641F^ is oncogenic and cooperates with *Braf*^V600E^ mutations and *Pten* loss to accelerate melanoma formation ^12^. Furthermore, we found that mutant *Ezh2*^Y641F^ co-immunoprecipitates with Stat3, and together they activate expression of several common target genes. One class of genes co-regulated by Ezh2 and Stat3 in *Ezh2*^Y641F^ mutant melanomas were MHC class I antigen processing genes in the H2-Q cluster, which are directly implicated in anti-tumor immunity ^18^. In addition to these MHC class I genes, chromatin immunoprecipitation followed by sequencing (ChIP-seq), suggests that Ezh2 and Stat3 are also found at the same promoter regions of the autophagy regulator, Atg7. Atg7 is a critical protein for autophagy initiation, as it facilitates an intermediate step in LC3 lipidation through its E1-like enzymatic activity ^19^. Atg7 conjugates with and adenylates LC3 (a ubiquitin-like protein also known as Atg8) and then transfers LC3 to the E2-like enzyme Atg3, which catalyzes the conjugation of LC3 to phosphatidylethanolamine (PE) on the autophagosome membrane ^20,21^. LC3 lipidation and, therefore, Atg7 are necessary for normal autophagosome formation, and Atg7 deficient cells are also autophagy deficient ^19,22^. Autophagy plays a significant role in many different cellular functions, both cell-intrinsically and extrinsically. In cancer, numerous autophagy regulators are mutated or deregulated ^23–25^, but given its role in many cellular mechanisms, its contribution during different phases of carcinogenesis is not entirely understood. In melanoma, previous studies have shown that deletion of *Atg7* in a mouse model driven by the oncogenic *Braf*^V600E^ and deletion of the tumor suppressor *Pten* significantly slowed down melanoma growth, suggesting that *Atg7* functions as an oncogene ^26^. Mechanistically, the study showed that deletion of *Atg7* resulted in increased oxidative stress and cellular senescence, which served as a barrier to melanomagenesis ^26^. Carcinogenesis, however, involves many different steps, from initial melanocyte transformation and immortalization to angiogenesis and immune evasion. The latter is particularly important in melanoma since checkpoint inhibitors have dramatically increased melanoma survival in the last ten years ^27–30^. Despite this improvement, many patients do not respond to treatment or experience severe toxicity, necessitating better understanding of anti-tumor immune mechanisms. Many autophagy components have been implicated in tumor immunity in multiple solid tumors ^31–35^, partially driven by their role in recycling unwanted cellular components and processing peptides, and may therefore play an important role in immunotherapy approaches.

Given our prior findings that *Ezh2*^Y641F^ mutant melanomas have a significantly altered tumor immunity, and the fact that Ezh2 and Stat3 can both be found at the *Atg7* locus, we hypothesized that *Atg7* may contribute to the altered tumor immune response in *Ezh2*^Y641F^ melanomas. In this study we investigated the role of Atg7 in both *Ezh2*^WT^ and *Ezh2*^Y641F^ melanoma tumor growth and its effect on anti-tumor immunity.

## RESULTS

### Ezh2 and Stat3 regulate Atg7 expression in melanoma cells

Previously, we investigated the role of *Ezh2*^Y641F^ mutations in melanoma and found a direct interaction of *Ezh2*^Y641F^ with Stat3, with direct effects on tumor immunity ^18^. We also identified a number of loci directly bound by both Ezh2 and Stat3 in melanoma cells. Here, we expanded that study to additional cell lines to gain a more comprehensive understanding of genes regulated by both Ezh2 and Stat3 in melanoma in a *Braf*^V600E^/Pten^F/F^ background, with or without the *Ezh2*^Y641F^ mutation. First, using Stat3 ChIP-seq, we confirmed enrichment of Stat3 binding motifs in *Ezh2*^Y641F^ melanoma cells compared to *Ezh2*^WT^ cells, and also identified enriched representation of motifs of other immune regulators, such as Stat1 and Irf1 (**Fig. 1a**). Gene Set Enrichment Analysis (GSEA) ^36^ of Stat3 peaks enriched in *Ezh2*^Y641F^ mutant melanoma cells identified several oncogenic signatures. Interestingly, we also identified several gene expression signatures that implicate autophagy or related cellular processes (**Fig. 1b**). We next assessed whether autophagy regulators were differentially expressed in *Ezh2*^WT^ vs *Ezh2*^Y641F^ melanoma ^12^. We found that *Atg7*, an important autophagy regulator, was upregulated in *Ezh2*^Y641F^ melanomas compared to *Ezh2*^WT^, and its expression was downregulated upon treatment with a pharmacological Ezh2 inhibitor (**Fig. 1c**). We also found increased expression of Atg7 in *Ezh2*^Y641F^ melanoma cells at the protein level (**Fig. 1d**). Chromatin immunoprecipitation followed by sequencing (ChIP-seq) analysis identified several Stat3 and Ezh2 peaks at the *Atg7* gene promoter and first intron (**Fig. 1f**). To confirm the relevance of these data in human patients, we analyzed data from the ReMap Atlas of regulatory regions (a collection of all public ChIP-seq data for transcriptional regulators from GEO, ArrayExpress, and ENCODE databases) ^37^ for EZH2 and STAT3 in various cell types and the ENCODE registry of candidate cis-regulatory elements^38^. We identified several cis-regulatory elements that coincide with mouse experimental Ezh2 and Stat3 binding sites (**Fig. 1e**), suggesting that our findings in mouse models are conserved and potentially relevant to human disease.

**Figure 1.**
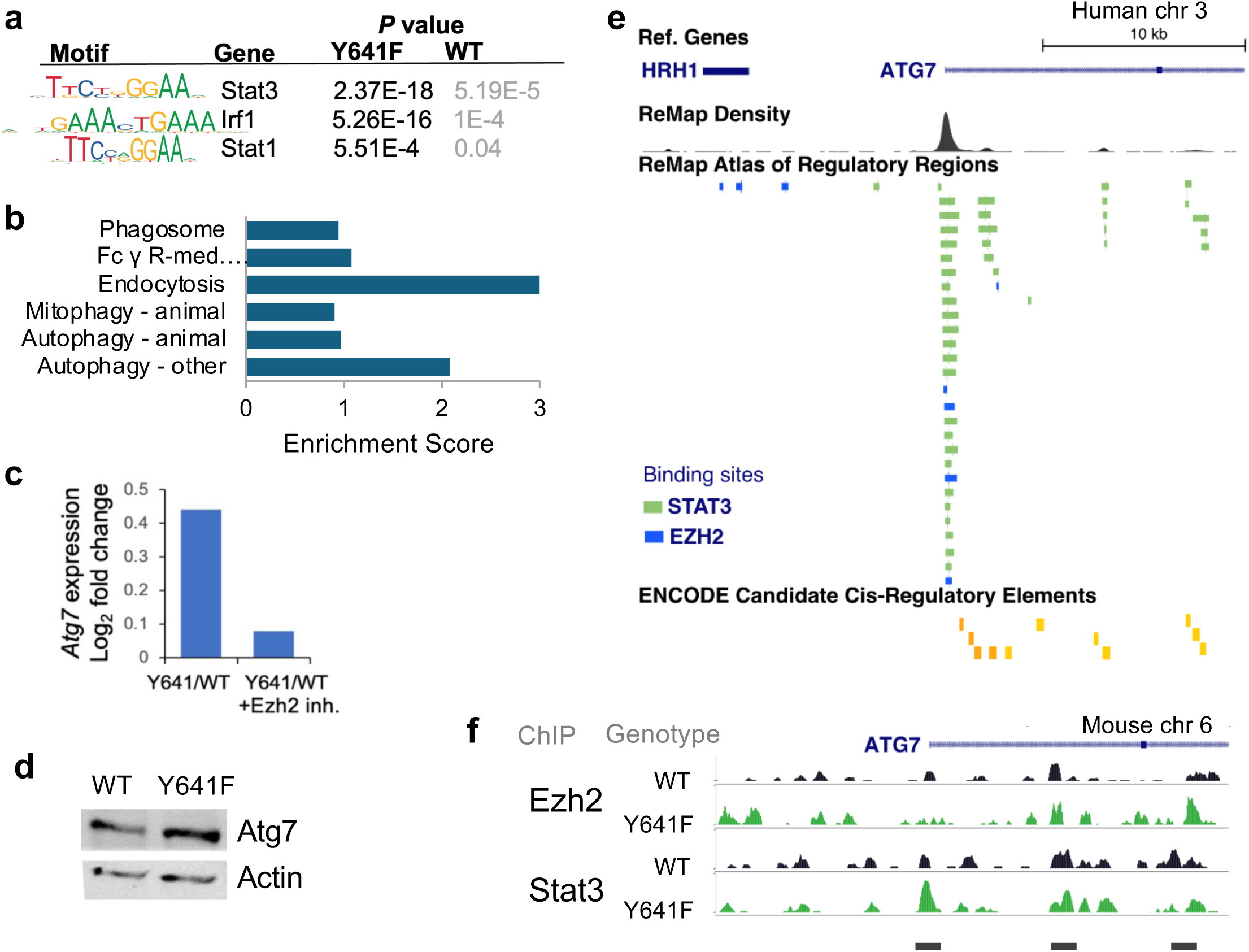
Regulation of *Atg7* expression by Ezh2 and Stat3. **(a)** Enriched motifs in *Ezh2*^WT^ and *Ezh2*^Y641F^ melanoma cells. **(b)** Gene Set Enrichment Analysis (GSEA) of Stat3 ChIP-seq peaks identifies several signatures associated with autophagy mechanisms (FDR<0.05). **(c)** Transcript expression of *Atg7* in *Ezh2*^Y641F^ vs *Ezh2*^WT^ melanoma cells, in the absence or presence of the Ezh2 inhibitor JQEZ5. **(d)** Protein expression of Atg7 in *Ezh2*^WT^ vs *Ezh2*^Y641F^ melanoma cells by Western blot. **(e)** Human ChIP-seq data in various cell lines showing direct binding of both STAT3 (green) and EZH2 (blue) at the *ATG7* promoter and intronic regions that correspond to cis-regulatory elements. Image modified from UCSC Genome Browser. **(f)** ChIP-seq tracks for Ezh2 and Stat3 in *Ezh2*^WT^ and *Ezh2*^Y641F^ melanoma cells at the mouse *Atg7* locus indicating binding at the *Atg7* promoter and first intron.

### Loss of Atg7 inhibits in vitro *and* in vivo cell growth

To determine whether Stat3 controls expression of Atg7, we generated stable Stat3 knockdown melanoma cells lines using shRNA. We found that *Stat3* knockdown in at least two independent mouse melanoma cell lines resulted in lower Atg7 protein levels, consistent with the hypothesis that Stat3 positively regulates *Atg7* expression (**Fig. 2a-c**). Since Atg7 is an important regulator of autophagy initiation, we assessed the ratio of type I cytosolic LC3 (LC3-I) and the type II lipid-conjugated form that is present on autophagosome membranes (LC3-II), a standard assay for assessing autophagy ^39,40^. We found that after Stat3 knockdown, cells exhibited a lower LC3-II/I ratio, indicating reduced levels of autophagy (**Fig. 2c**), consistent with depletion of Atg7 protein levels. We next investigated whether Atg7 is required for *in vitro* melanoma growth. We used a lentiviral CRISPR/Cas9 system to inactivate *Atg7* expression in two *Ezh2*^WT^ and two *Ezh2*^Y641F^ melanoma cell lines. The lentiviral system is a single vector delivery of the single guide RNA (sgRNA) targeting Atg7, Cas9, puromycin for selection and GFP for cell sorting ^41^. For controls we generated stable cell lines using two non-specific sgRNAs. After puromycin selection, GFP+ transfected cells were sorted by FACS and tested for knockout efficiency by Western blot. We identified multiple clones that exhibited small genomic deletions within the targeted exon and which resulted in complete loss of Atg7 protein expression (**Fig. 2d-e**). We further tested these clones for autophagy activity, and they exhibited a decreased LC3-II/I ratio, verifying disruption of Atg7 function and lower autophagic activity (n=4, p<0.05) (**Fig. 2d-e**). To determine whether absence of Atg7 affects cell intrinsic melanoma growth *in vitro*, we monitored cell growth using Alamar Blue staining, a cell-permeable dye, (resazurin), which serves as a redox indicator in response to cellular metabolic activity ^42^. We found that deletion of *Atg7* only transiently slowed the growth of *Ezh2*^WT^ cells, but did not have a significant overall effect during the duration of the *in vitro* assay (**Fig. 2f**), or an effect on the growth rate of *Ezh2*^Y641F^ melanoma cells. These results suggest that the effect of *Atg7* deletion on melanoma cell growth may depend not only on increased cellular stress and senescence, as previously suggested ^26^, but also on specific *in vivo* variables and cell extrinsic factors such as the tumor microenvironment and anti-tumor immunity.

**Figure 2.**
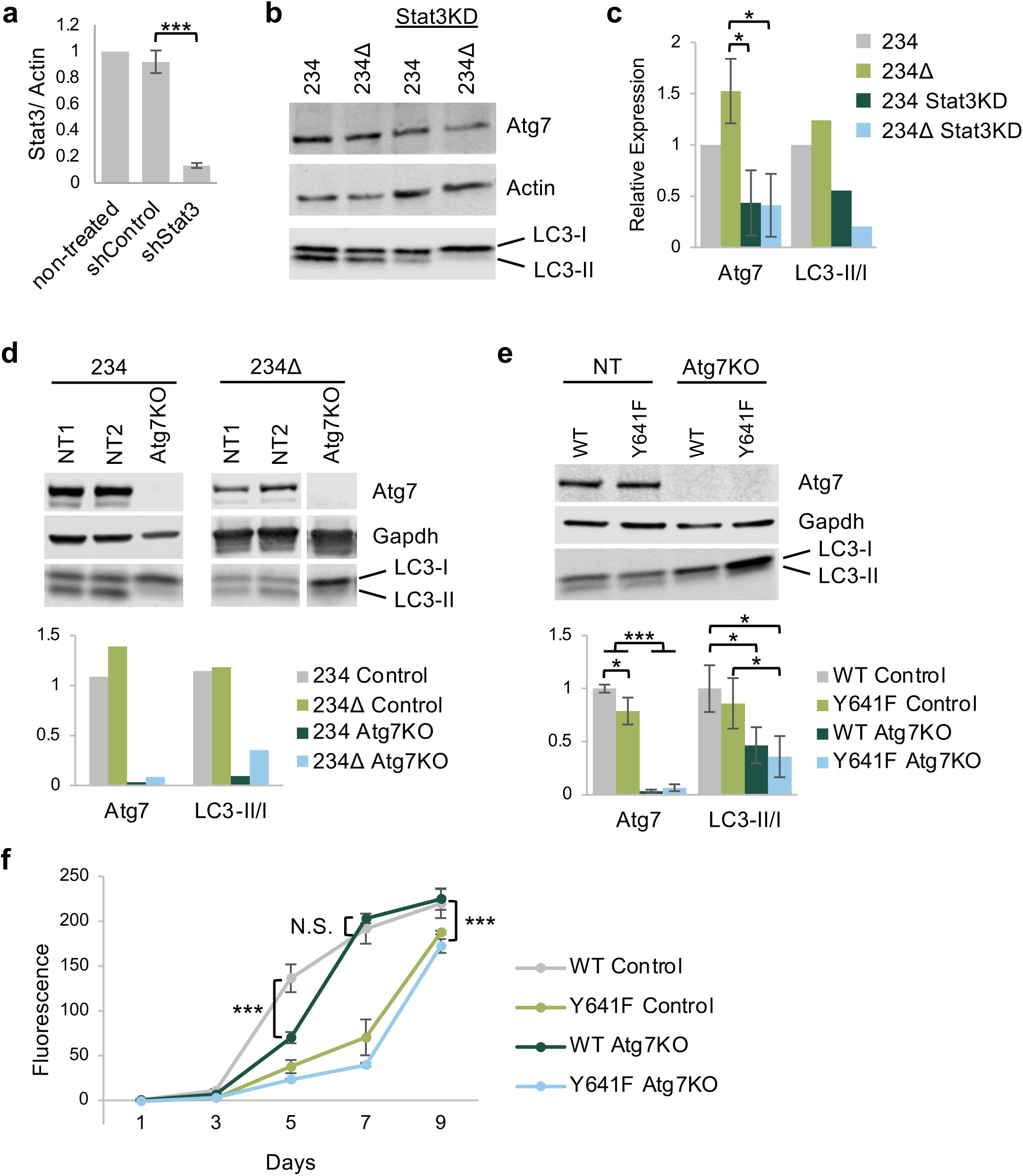
Deletion of *Atg7* in melanoma cells has no significant effect on cell intrinsic cell growth *in vitro*. **(a)** Protein expression of Stat3 after shRNA-mediated stable gene knockdown in melanoma cell line 234Δ. **(b)** Expression of Atg7 and LC3 after Stat3 knockdown in *Ezh2*^WT^ and *Ezh2*^Y641F^ melanoma cell lines 234 and 234Δ. **(c)** Quantification of western blot in b. Atg7/Actin N = 2, LC3-II/I N = 1. **(d)** (Top) Immunoblotting for Atg7 and LC3-I/II in control and *Atg7* knockout clones in the 234 and 234Δ cell lines. NT, non-targeted sgRNA. (Bottom) Quantification of the plots above, Atg7/Gapdh and LC3-II/I N = 1. **(e)** As in (d), with a second set of *Ezh2*^WT^ (27.6-M2) and *Ezh2*^Y641F^ (28.2-M4) melanoma cell lines. Atg7/Gapdh N = 3, LC3-II/I N = 4. **(f)** *In vitro* growth curve of *Ezh2*^WT^ and *Ezh2*^Y641F^ melanoma cell lines 27.6-M2 and 28.2-M4 with and without *Atg7* deletion. N.S. = not statistically significant. For all graphs error bars are standard deviation, *** p-value <0.001, * p-value <0.05.

### *Atg7* deletion suppresses *in vivo* tumor growth and results in increased CD8+ T cells in the tumor microenvironment

To test whether Atg7 deletion differentially affects *in vivo* growth of *Ezh2*^WT^ or *Ezh2*^Y641F^ mutant melanomas, we adaptively transferred five hundred thousand *Atg7* knockout *Ezh2*^WT^ and *Ezh2*^Y641F^ melanoma cells into the left and right flank of wildtype recipient mice. These cells formed tumors, which we monitored for growth over time. Consistent with our prior finding, tumors expressing *Ezh2*^Y641F^ grew more slowly than *Ezh2*^WT^ ^18^, and deletion of *Atg7* resulted in slower tumor growth regardless of *Ezh2* status (n=8, p<0.001 for WT Control vs all other groups at every time point) (**Fig. 3a**). These results are consistent with a prior study that demonstrated the oncogenic activity of *Atg7* in a *Braf*^V600E^/*Pten*^F/F^ background ^26^, which was attributed to a cell intrinsic increase of oxidative stress and senescence of the tumor cells, without consideration of cell extrinsic variables. Since we previously showed that tumor immunity is an important factor in the progression of *Ezh2*^Y641F^ melanomas *in vivo*, we investigated how deletion of *Atg7* affected infiltration of immune cells in *Ezh2*^WT^ and *Ezh2*^Y641F^ melanomas. We harvested tumors seven days after injection and analyzed tumor immune cell infiltration by flow cytometry. We found that the overall amount of CD45+ tumor infiltrating cells, while somewhat variable, tended to be higher after *Atg7* deletion, particularly in *Ezh2*^WT^ melanoma tumors (n=8, p=0.024) (**Fig. 3b**). Nevertheless, we observed more significant differences in the type of immune cells that infiltrated these tumors. In the *Ezh2*^Y641F^ control group, we detected increased CD8+ T cell infiltration compared to *Ezh2*^WT^ (n=8, p<0.001), confirming our prior findings ^18^. Deletion of *Atg7* resulted in no change to CD8+ T cell infiltration in *Ezh2*^WT^; however, *Atg7* deletion in *Ezh2*^Y641F^ tumors resulted in an approximately 2-fold increase in the CD8+ population (n=7-8, p<0.001) (**Fig. 3c-d**). Interestingly, we found that expression of *Ezh2*^Y641F^, regardless of *Atg7* expression, dramatically increased infiltration of natural killer (NK) cells, a population that we had not previously assessed in this model (n=7-8, p<0.001) (**Fig. 3c**). Other lymphoid populations such as CD4+ cells were elevated in *Ezh2*^Y641F^ compared to *Ezh2*^WT^ tumors, but deletion of *Atg7* had no significant effect compared to the control group in either *Ezh2* genotype (**Fig. 3c-d**).

**Figure 3.**
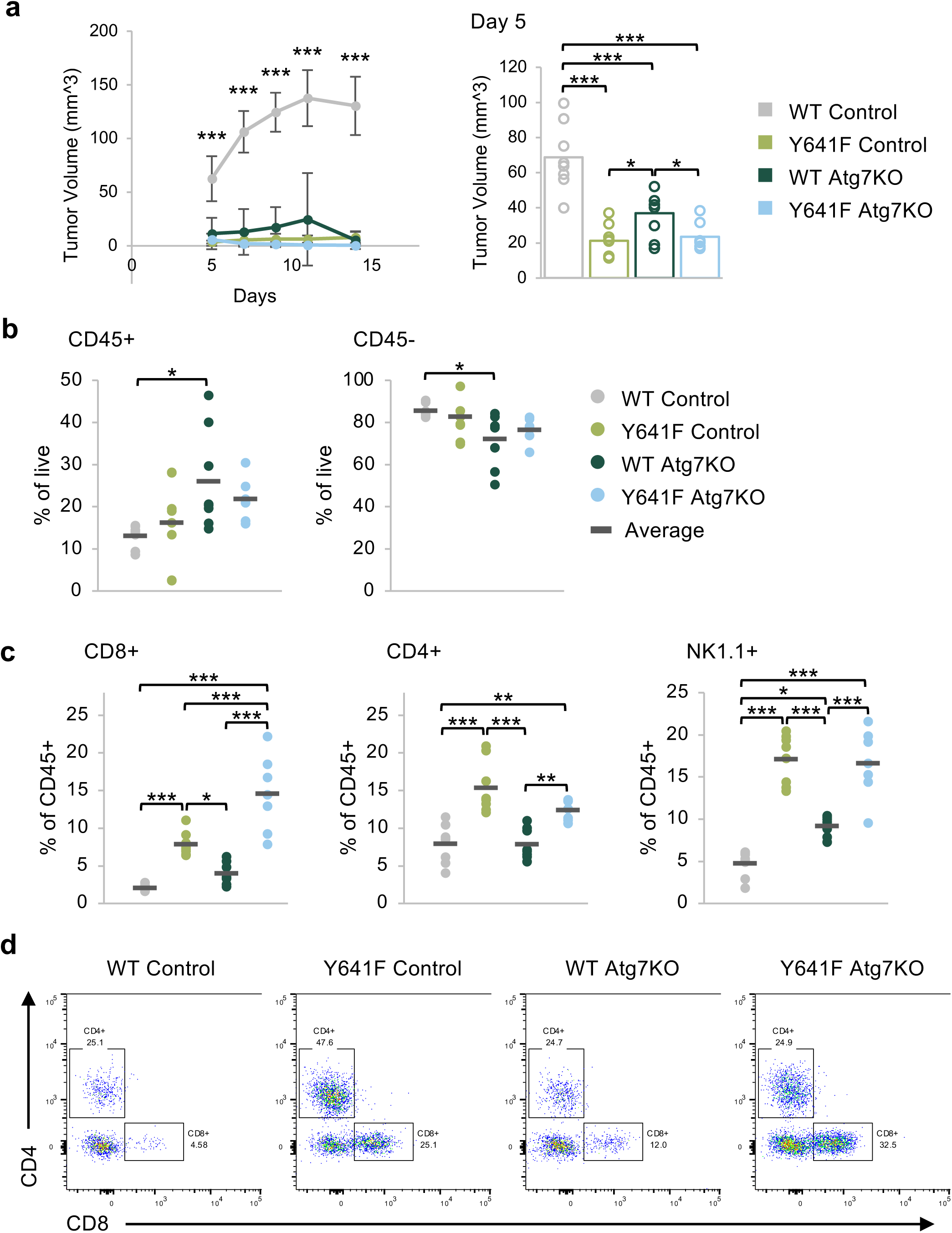
Deletion of *Atg7* in melanoma cells results in slower *in vivo* tumor growth and increased presence of tumor infiltration of lymphocytes. **(a)** (Left) *In vivo* tumor growth in *Ezh2*^WT^ (27.6M2) and *Ezh2*^Y641F^ (28.2M4) melanomas, with and without *Atg7* deletion. The group average is displayed and error bars indicate the standard deviation. Control = non-targeted sgRNA, N = 8 per group, representative of two independent experiments. (Right) Tumor volume at day 5 post injection. The bars indicate the group mean, and the circles are individual tumors. **(b)** Flow cytometric analysis of tumor infiltrating CD45+ hematopoietic cells and CD45-cells. N = 6-8 tumors per group. **(c)** Flow cytometric analysis of tumor infiltrating CD8+, CD4+ and NK1.1+ cells. N = 7-8 tumors per group. **(d)** Representative flow cytometry plots of the CD4+ and CD8+ data shown in panel c. For the graphs in b and d, each dot on the graph represents an individual tumor, and the black bar marks the average for the group. *p<0.05, **p<0.01, ***p<0.001.

While the number of cytotoxic CD8+ T cells significantly increased with *Atg7* deletion in *Ezh2*^Y641F^ melanoma, it is possible that these T cells are not functionally competent killer cells. T cells have evolved mechanisms to prevent autoreactivity through receptor-ligand interactions, also known as immune checkpoints. These interactions are very important in cancer, as ligands expressed on tumors may interact with receptors on T cells to inhibit anti-tumor activity. One such immune checkpoint pair is PD-1 and PD-L1. We thus assessed the presence of the PD-1 on T cells in the tumor microenvironment, and PD-L1 on the melanoma cells. We found increased expression of PD-1 in CD8+ T cells after *Atg7* knockout (n=7-8, p<0.001) and to a lesser degree in CD4+ cells (**Fig. 4a**). *Ezh2*^Y641F^ *Atg7* knockout tumors also exhibited increased expression of PD-L1 compared to all other groups (p<0.05) (**Fig. 4b**). These data suggest that while loss of *Atg7* results in slower tumor growth, likely partially mediated by increased presence of CD8+ T cells, it may also eventually lead to T cell exhaustion.

**Figure 4.**
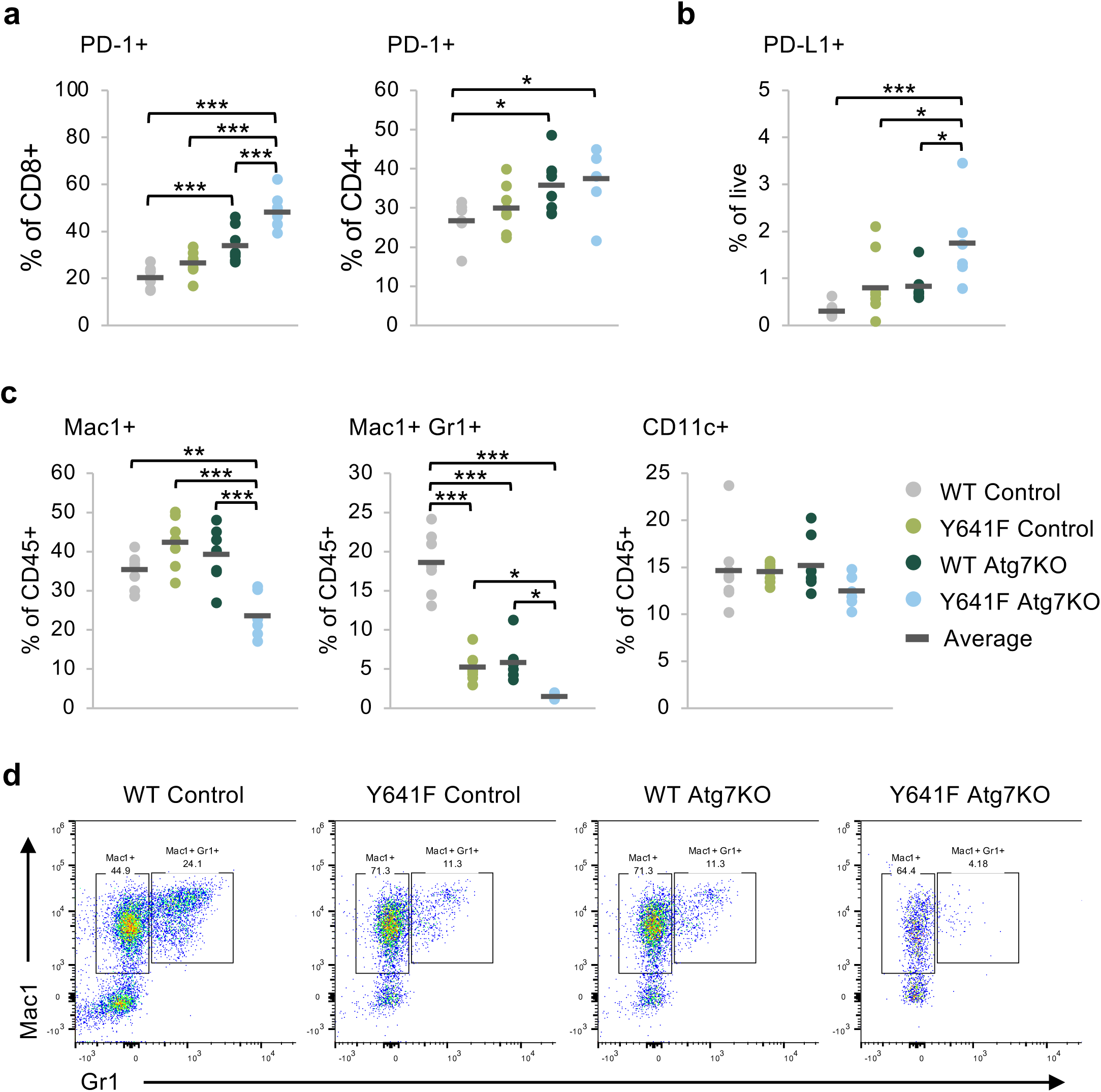
Deletion of *Atg7* in melanoma cells results in decreased infiltration of myelosuppressive cells. **(a)** Expression of PD1 on tumor infiltrating CD8+ and CD4+ T cells in *Ezh2*^WT^ and *Ezh2*^Y641F^ melanoma cells, with and without *Atg7* deletion. N = 7-8 tumors per group. **(b)** Expression of the PD1 ligand (PD-L1) on the melanoma cells from panel a. N = 6-8 tumors per group. **(c)** Flow cytometric analysis of tumor infiltrated CD11c+, Mac1+ and double Mac1/Gr1+ cells in *Ezh2*^WT^ and *Ezh2*^Y641F^ melanoma tumors, with and without *Atg7* deletion. N = 6-8 tumors per group. **(d)** Representative flow cytometry plots for the Mac1+ and double Mac1/Gr1+ data in panel c. For the graphs in a-c, each dot on the graph represents an individual tumor, and the black bar marks the average for the group. *p<0.05, **p<0.01, ***p<0.001.

### Deletion of *Atg7* results in a decrease of myelosuppressive cells in the melanoma tumor microenvironment

Another important immune population that plays a critical role in tumor immunity are myeloid-derived suppressor cells (MDSCs). To test whether *Atg7* deletion affects infiltration of these cells in the melanoma tumor microenvironment, we measured expression of myeloid markers using flow cytometry. We found a significant decrease of Mac1+/Gr1+ double-positive cells after *Atg7* deletion in both *Ezh2*^WT^ and *Ezh2*^Y641F^ cells (n=6-8, p<0.001 WT, p<0.05 Y641F), with a significantly lower frequency in the *Ezh2*^Y641F^ tumors (p=0.04), while Mac1+ cells decreased only in the *Ezh2*^Y641F^ *Atg7* knockout tumors (n=6-8, p<0.01) (**Fig. 4c-d**). We did not find changes in the dendritic cell population as determined by CD11c expression in any of the groups, regardless of *Ezh2* status or *Atg7* expression (**Fig. 4c**).

Overall, these results suggest that deletion of Atg7 affects the recruitment of both lymphoid and myeloid populations in the melanoma tumor microenvironment. Some of the observed phenotypes were stronger when *Atg7* was deleted in the presence of *Ezh2*^Y641F^, suggesting that some of the effects of *Ezh2*^Y641F^ on melanoma tumor immunity may be mediated by Atg7. It remains to be seen whether the effects of Atg7 on tumor immunity are mediated through its role in autophagy or whether they are mediated by autophagy-independent, cell intrinsic mechanisms.

## DISCUSSION

In this study we investigated the role of downstream targets of Ezh2 in the melanoma tumor immune response. Ezh2 regulates many different hallmarks of cancer, from cell intrinsic cell cycle regulation to tumor immunity. Ezh2 has a complex role in cancer. It is often deleted in some cancers while amplified in others, consequently functioning both as a tumor suppressor and as an oncogene. While typically functioning within the PRC2 complex and mediating methylation of lysine 27 on histone 3, Ezh2 can also function independently of the PRC2 complex, sometimes as a transcriptional activator as we and others have previously shown ^18,43^. Here we investigated the role of one of its non-canonical targets, Atg7, an autophagy regulator.

Autophagy is a fundamental cellular mechanism required to maintain cellular health. When perturbed it can result in the onset of different diseases. In antigen-presenting cells, such as dendritic cells, autophagy generates peptides from endogenous antigens, which are presented by MHC class II proteins to CD4+ cells to prime the immune response. In cancer, the role of autophagy is context dependent. Autophagy in tumor cells can enhance processing of exogenous antigens and MHC-I antigen presentation, inducing CD8 T cell priming and cytotoxic activity ^44^. Specifically, *ATG* genes, such as *ATG7*, are involved in the internalization and recycling of the MHC-I molecules themselves ^44^, and dendritic cells deficient in Atg7 have increased cell surface expression of MHC-I molecules ^45^. Autophagy, therefore, can stimulate CD8+ T cells, thus functioning in a tumor suppressive manner ^46^. In our melanoma models, it is possible that deletion of *Atg7* similarly increases the amount of MHC-I at the cell surface, resulting in the increased CD8+ T cell infiltration that we observe in melanoma tumors. On the other hand, because cancer cells require autophagy for growth, autophagy-regulating genes can also function as oncogenes ^26^. Consistent with an oncogenic function, in humans, melanoma patients with a high autophagic index benefit less from chemotherapy, exhibit increased tumor cell proliferation and metastasis, and have poor outcomes ^47,48^. Overall, this dual role of autophagy in cancer is not well understood and may be context dependent.

Within the context of *Ezh2*^Y641F^-mutant melanomas, loss of *Atg7* does not have a significant effect on cell intrinsic cell growth, but it appears to further enhance anti-tumor immunity with increased presence of cytotoxic CD8+ T cells and decreased MDSCs populations in the tumor microenvironment, a combination that is not conducive to tumor growth. Expression of *Atg7* does not change dramatically with expression of *Ezh2*^Y641F^, *in vitro*, but its expression is regulated by Stat3, as clearly demonstrated with Stat3 knock-down experiments. Ezh2 and Stat3 may, therefore, play a role in sustaining *Atg7* expression within the context of a more complicated transcriptional network, and that Atg7 may be playing a secondary role in the many role of *Ezh2*^Y641F^ mutations in melanoma.

Tumor immunobiology is very complex and is affected by a multitude of factors, including cell-intrinsic variables, the stroma, fibrosis, the tumor tissue location, the vasculature, tumor burden, signals or cytokines secreted by tumor cells, and others. It is possible that deletion of Atg7 affects any of these factors, whether via autophagy-dependent or -independent functions. Regardless of the mechanisms, our results indicate the relevance of tumor immunity in melanoma tumors lacking expression of *Atg7*. Future studies are needed to further delineate mechanistically how *Atg7* deletion results in such significant changes to the tumor immune response in melanoma and how it cooperates with mutations in Ezh2. With the availability of several pharmacological inhibitors of autophagy mechanisms, our study suggests that targeting autophagy-related pathways could be a viable strategy to modulate anti-tumor immunity, offering potential for therapeutic advancements in melanoma treatment.

## MATERIALS & METHODS

### Genomic analysis

ChIP-seq and RNA-seq were performed on *Ezh2*^WT^ and *Ezh2*^Y641F^ mouse melanoma cells with or without treatment with the Ezh2 inhibitor JQEZ5 as described previously ^18^. Analysis of transcription factor motif enrichment was carried out using HOMER ^49^. Functional significance of Ezh2 and Stat3 binding sites/peaks was evaluated using the Genomic Regions Enrichment of Annotations Tool (GREAT) ^50^ and Gene Set Enrichment Analysis was performed as described here ^36^. The UCSC Genome Browser was used to visualize EZH2 and STAT3 binding sites at the ATG7 locus (human GRCh38/hg38) using tracks for the ReMap Atlas of Regulatory Regions and the ENCODE Candidate Cis-Regulatory Elements (cCREs) ^37^.

### Cell culture & CRISPR knockouts

Eight mouse melanoma cell lines were used: 234, 480, and 855 (*Ezh2*^WT^ *Tyr-CRE*^ERT2^ *Braf*^V600E/+^ *Pten*^flox/flox^); 234Δ, 480Δ, and 855Δ (*Ezh2*^Y641F^ *Tyr-CRE*^ERT2^ *Braf*^V600E/+^ *Pten*^flox/flox^); 27.6-M2 (*Ezh2*^WT^ *Tyr-CRE*^ERT2^ *Braf*^V600E/+^ *Pten*^flox/+^); and 28.2-M4 (*Ezh2*^Y641F^ *Tyr-CRE*^ERT2^ *Braf*^V600E/+^ *Pten*^flox/+^). Cells were cultured in DMEM (Sigma D6429) with 10% FBS (Corning Cat# MT35010CV) and 1% penicillin-streptomycin (Genesee Scientific Cat# 25-512). *Atg7* knockout cell lines were generated by transducing cells with lentiviral CRISPR/Cas9 (TLCV2 Addgene plasmid #87360). Lentiviruses were generated using 293T cells via transfection with PEI. Stable cell lines were selected by treating with puromycin for 7 days (3 µg/ml, refreshed every other day), and Cas9 expression was induced with 3-5 doses of doxycycline at 1 µg/ml. To generate single clones, GFP+ and PI negative cells were single cell sorted into 96 well plates on the MoFlo sorter (Beckman Coulter) at the Siteman Flow Cytometry Core Facility. The clones were tested for *Atg7* knockout by immunoblotting. For the in vitro cell growth assay, cells were plated at 1000 cells/well in a 24-well plate in triplicate, one set of triplicates for each time point. For each measurement, the growth media was aspirated and replaced with media containing Alamar Blue (Invitrogen #A50100) cell viability reagent at 1:10 dilution ^42^. The cells were returned to the incubator for 1 hour, after which 100 µl of supernatant was transferred from the 24-well plate to a clean 96-well plate. The samples were scanned on a BioTek Synergy HT plate reader using fluorescent excitation at 485/20 nm and detection at 590/35 nm. Data analysis was performed in Excel and statistically significant differences were determined by one-way ANOVA.

### Immunoblotting

Samples were prepared in Laemmli buffer with beta-mercaptoethanol, run on 4-20% pre-cast gels (Bio-Rad Mini-PROTEAN TGX Gels Cat# 4561095) using the BioRad Mini-PROTEAN system, and then transferred onto nitrocellulose membranes. The membranes were blocked for 1 hour in 5% milk in TBS-T, and then incubated with primary antibodies overnight at 4°C. Primary antibodies: anti-ATG7 (Cell Signaling #8558 at 1:500), anti-ACTIN (Abcam ab213262 at 1:1000), anti-GAPDH (Cell Signaling #5174 at 1:1000), and anti-LC3A/B (Cell Signaling #12741 at 1:1000). Membranes were washed with TBS-T before staining with secondary anti-rabbit IgG (H+L) DyLight 800 4X PEG Conjugate (Cell Signaling #5151) at 1:20,000 at room temperature for 1 hour. Membranes were imaged using a Licor Odyssey Infrared Imager, and Image Studio software was used for densitometry analysis. Statistically significant differences were detected using one-way ANOVA.

### Animals

Animals were housed in an Association for Assessment and Accreditation of Laboratory Animal Care (AAALAC)-accredited facility and treated in accordance with protocols approved by the Institutional Animal Care and Use Committee (IACUC) for animal research at Washington University in St. Louis.

### *In vivo* tumor models

Wildtype C57Bl/6 mice were generated in house. Tumor cells in Matrigel (Corning 354234) were injected subcutaneously in the flank at 0.5 × 10^6 cells per injection, two injections per mouse. Tumor growth was measured using digital calipers on day 5 post-injection and then every other day. For the flow cytometry analysis of tumor-infiltrating lymphocytes, tumors were harvested at day 7. Tumors were chopped in HBSS, dispersed using a syringe with 18G needle, and passed through a 0.40 µm filter.

### Flow Cytometric analysis

Single cell suspensions from tumors were washed with HBSS containing 2% FBS and 1 mM EDTA and stained with the following antibody cocktails for detecting lymphoid populations: anti-CD45-PerCP/Cy5.5 (BioLegend 103132), anti-NK1.1-FITC (BioLegend 108706), anti-CD3-PB (BioLegend 100214), anti-CD4-APC (BioLegend 100412), anti-CD8-AF700 (BioLegend 100730), and anti-PD-1(CD279)-PE/Cy7 (BioLegend 135216), and myeloid populations: anti-CD45-PerCP/Cy5.5 (BioLegend 103132), anti-CD19-FITC (BioLegend 115506), anti-B220-FITC (BioLegend 103206), anti-CD3-FITC (BioLegend 100204), anti-CD11b(Mac1)-PB (BioLegend 101224), anti-CD11c-PE/Cy7 (BioLegend 117318), and anti-Ly-6G(Gr1)-AF700 (BioLegend 127622). Propidium Iodide was used to exclude dead cells. Samples were run on an Attune Nxt Flow Cytometer (ThermoFisher Scientific) at the Siteman Flow Cytometry Core Facility, analysis was done in FlowJo V10, and statistically significant differences were identified using One-way ANOVA.

## ACKNOWLEDGEMENTS

We thank the Siteman Flow Cytometry facility and the Department of Comparative Medicine for animal expertise. We also thank all members of the Souroullas lab for critical input on the manuscript. This work was supported by the Alvin J. Siteman Cancer Center, The Harry J. Lloyd Charitable Trust (GPS) and T32 CA113275-10 (SZ),

## AUTHOR CONTRIBUTIONS

GPS and SMZ designed experiments and wrote the manuscript. GPS, SMZ, ES and SS performed experiments, analyzed, and interpreted the data. ES and SS performed experiments. GPS conceived of and supervised the study.

## CONFLIC OF INTEREST

The authors declare no relevant competing financial interests.

